# SARS-CoV-2 accessory protein ORF8 decreases antibody-dependent cellular cytotoxicity

**DOI:** 10.1101/2022.03.30.486403

**Authors:** Guillaume Beaudoin-Bussières, Ariana Arduini, Catherine Bourassa, Halima Medjahed, Gabrielle Gendron-Lepage, Jonathan Richard, Qinghua Pan, Zhen Wang, Chen Liang, Andrés Finzi

## Abstract

SARS-CoV-2 Spike glycoprotein is the major target of host neutralizing antibodies and the most changing viral protein in the continuously emerging SARS-CoV-2 variants as a result of frequent viral evasion from host antibody responses. In addition, SARS-CoV-2 encodes multiple accessory proteins that modulate host antiviral immunity by different mechanisms. Among all SARS-CoV-2 accessory proteins, ORF8 is rapidly evolving and a deletion in this protein has been linked to milder disease. Here, we studied the effect of ORF8 on peripheral blood mononuclear cells (PBMC). Specifically, we found that ORF8 can bind monocytes as well as NK cells. Strikingly, ORF8 binds CD16a (FcγRIIIA) with nanomolar affinity and decreases the overall level of CD16 at the surface of monocytes and, to a lesser extent, NK cells. Strikingly, this decrease significantly reduces the capacity of PBMCs and particularly monocytes to mediate antibody-dependent cellular cytotoxicity (ADCC). Overall, our data identifies a new immune-evasion activity used by SARS-CoV-2 to escape humoral responses.

## Text

Since the discovery of the Severe Acute Respiratory Syndrome Coronavirus 2 (SARS-CoV-2) in early 2020 in Wuhan, China (1–3), major advances have been made in understanding this virus and the disease it causes. A large part of these advances have focused on one SARS-CoV-2 protein, its Spike, which resulted in the development of Spike therapeutic antibodies that reached the clinic and currently-approved vaccines. However, SARS-CoV-2 has a large and complex genome coding for 26 proteins (4) which all play essential roles in viral replication and pathogenesis. Most of these viral proteins remain unfortunately poorly studied compared to the Spike glycoprotein. This include the small (121 amino acids) rapidly evolving accessory protein ORF8 (5). This protein has two dimerization interfaces (6) and has been linked to immune evasion. Notably, ORF8 was shown to directly interact with major histocompatibility complex class I molecules (MHC-I) and mediate their downregulation by targeting them to lysosomal degradation via autophagy rendering infected cells more resistant to lysis by cytotoxic T cells (7). Interestingly, a recent study has shown that ORF8 is secreted from infected cells, can be detected in the plasma of acutely-infected individuals, and is negatively associated with survival (8). Whether this association is linked to its capacity to induce a cytokine storm remains to be determined (9).

Vaccine-elicited humoral responses were shown to protect from infection and severe disease (10, 11). Anti-SARS-CoV-2 Spike-specific antibodies can mediate a wide range of actions from viral neutralization to different Fc-mediated effector functions. Among the later, antibody-dependent cellular cytotoxicity (ADCC) and antibody-dependent cellular phagocytosis (ADCP) result in the elimination of infected cells. In the transgenic human ACE2 K18 mice model, Fc-mediated effector functions of neutralizing antibodies have been shown to be required for protection from lethal challenges (12). Similarly, strong Fc-mediated effector functions, in the absence of neutralization, were sufficient to significantly delay death (13). Furthermore, Fc-mediated effector functions were associated with survival in acutely infected individuals (14, 15). Altogether, these results suggest that Fc-effector functions have a major impact on virus clearance and disease outcomes. Accordingly, Fc-effector functions were associated with protection from infection by emerging SARS-CoV-2 variants of concern (16). These documented inhibitory impacts of Fc-effector functions on virus replication and disease might explain why viruses have developed sophisticated strategies to evade this important immune function (17–20). For example, HIV-1 uses two accessory proteins, Nef and Vpu, to protect infected cells from ADCC responses (21–24). Other viruses, such as Herpes simplex virus type 1 (HSV-1) and type 2 (HSV-2), murine cytomegalovirus (MCMV), human cytomegalovirus (HCMV) and varicella-zoster virus (VZV) secrete proteins that bind to the Fc portion of host immunoglobulins (25–28). These virally encoded Fc binding proteins are thought to contribute to protection from Fc-effector functions. Since acute SARS-CoV-2 infection is associated with a loss of CD16+ cells, in the present study we evaluated whether soluble ORF8 could modulate Fc-effector functions.

We first evaluated whether ORF8 could interact with different cell types present in peripheral blood mononuclear cells (PBMCs). To investigate this possibility, recombinant ORF8 was conjugated with FITC and incubated on ice with PBMCs for 30 minutes. Ten percent of cells present in PBMCs were bound by ORF8 (Fig. 1A). By further investigating which cell type bound ORF8 using anti-CD14, anti-CD16 and anti-CD56 antibodies, we found that ORF8 bound monocytes (CD14+CD16+) as well as NK cells (CD56+CD16+) (Fig. 1A). Because it has been previously shown that ORF8 adopts an Ig-like fold (6) and because CD16 is a common receptor at the surface of these two cell types, we asked whether ORF8 could directly interact with CD16a. Using bio-layer interferometry (BLI), we measured the affinity of recombinant ORF8 with the CD16a ectodomain. As presented in figure 1B, ORF8 bound CD16a with nanomolar affinity (38.9 nM, Fig. 1B).

**Figure 1.**
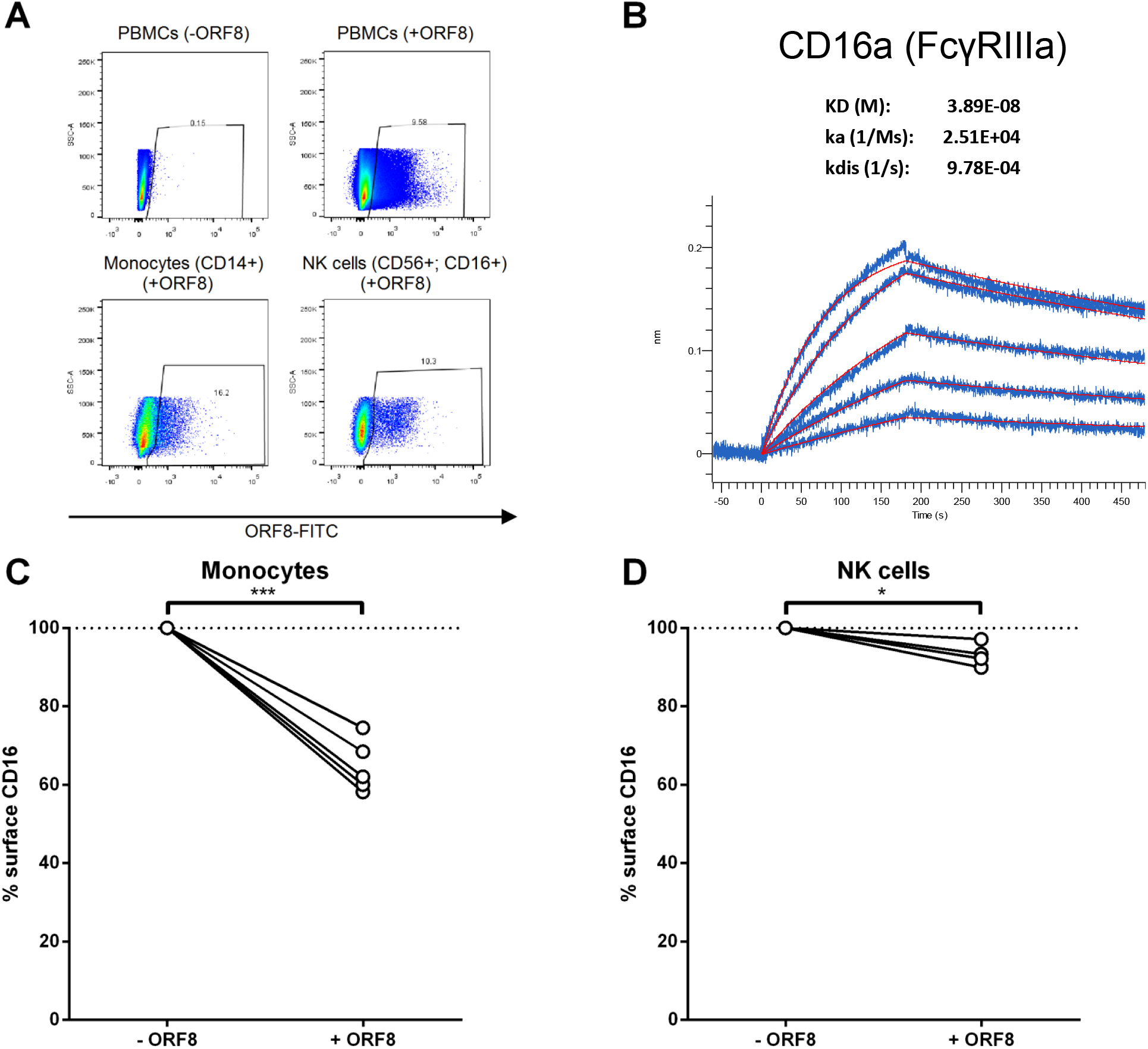
ORF8 binds monocytes and NK cells through CD16a. **(A)** PMBCs were incubated with FITC-conjugated recombinant ORF8 protein on ice for 30 min, followed by staining with anti-CD14-V450, and anti-CD16-PE-CY7 and anti-CD56-PE antibodies before being fixed with 4% PFA and analyzed with flow cytometer. **(B)** AR2G biosensors loaded with CD16a protein were soaked in two-fold dilution series of ORF8 (31.25 nM - 500 nM). Raw data are shown in *blue* and model in *red*. The affinity constant (KD), on rate (Ka) and off rates (Kdis) were calculated using a 1:1 binding model. (**C, D**) PBMCs from different donors were thawed and incubated for 16 hours with ORF8. The following day, the PBMCs were stained with anti-CD3, anti-CD14, anti-CD56, anti-CD16 and LIVE/DEAD Fixable Aqua Dead Cell Stain and analyzed by flow cytometry to measure surface levels of CD16 on (**C**) monocytes (n=5) and (**D**) NK cells (n=4). Cell-surface CD16 levels in presence of ORF8 were normalized on cell-surface CD16 detected in absence of ORF8. Statistical significance was evaluated using a parametric paired t-test.

CD16 interacts with the Fc portion of IgG and is present at the surface of different cell types including NK cells and monocytes. Upon binding a sufficient amount of antibodies, these cells get activated and can mediate ADCC. To evaluate whether ORF8 could modulate this response, we first assessed whether ORF8 modulated CD16 levels at the surface of NK cells and monocytes. Briefly, PBMCs were incubated overnight with ORF8 as described in material and methods. Sixteen hours after ORF8 treatment, PBMCs were stained with anti-CD3, anti-CD14, anti-CD56 and anti-CD16 antibodies and analyzed by flow cytometry to measure CD16 levels at the surface of NK cells and monocytes. As presented in figure 1C, the surface level of CD16 on monocytes present within the PBMC population was significantly decreased upon treatment with ORF8. When looking at CD16 at the surface of NK cells, a smaller, but significant, decrease was also observed (Fig. 1D). To evaluate if ORF8 directly acted on monocytes to decrease their CD16 level, we repeated the experiment using monocytes purified from PBMCs. As shown in supplemental figure 1A, addition of soluble ORF8 to purified monocytes modulated CD16 levels in a manner similar to when added to total PBMCs, therefore suggesting a direct effect of ORF8 on monocytes. To verify if the decreased CD16 levels at the surface of NK cells also depends on the presence of monocytes, ORF8 was added to monocyte-depleted PBMCs. As shown in supplemental figure 1B, the small decrease on NK cells was not observed upon monocyte depletion, confirming the role of monocytes in ORF8-mediated downmodulation of CD16 on NK cells. These results suggest that monocytes represent an important target of ORF8.

We next evaluated if the observed decrease of CD16 at the surface of monocytes and NK cells affected ADCC responses with *in vitro* assay which uses PBMCs as effector cells (29, 30). We first produced soluble ORF8 in culture media by transfecting HEK293T cells with ORF8 DNA. In agreement with previous data showing ORF8 secretion (8, 9), we also observed ORF8 secretion and supernatant accumulation (Fig. 2A). PBMCs were treated overnight with ORF8-conditioned media before being used as effector cells in the ADCC assay. In agreement with ORF8-mediated CD16 downregulation, soluble ORF8 significantly decreased ADCC responses mediated by plasma from convalescent and vaccinated individuals (Fig. 2B-D). The contribution of monocytes in these results was assessed using ORF8-treated monocytes. In agreement with decreased CD16 levels at the surface of ORF8-treated monocytes, we observed a significant decline in monocyte-mediated ADCC (Fig. 2E). Of note, the decrease in CD16 levels and ADCC responses after the addition of ORF8 to total PBMCs or purified monocytes was similar (Supplemental Figure 1 C and D), suggesting that the impact of ORF8 on PBMCs is mostly dependent on monocytes.

**Figure 2.**
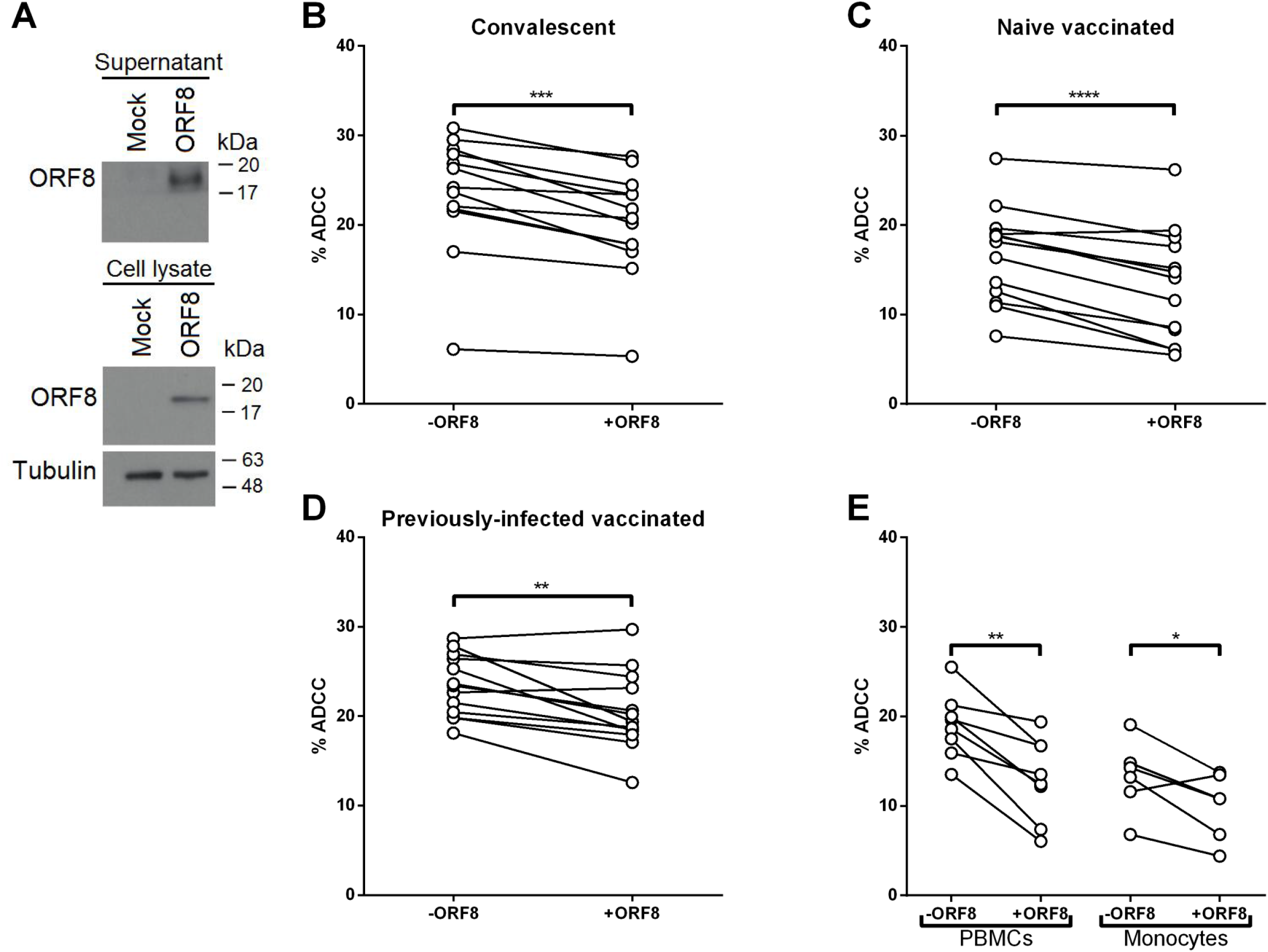
ORF8 decreases PBMC-mediated ADCC. (**A**) Secretion of ORF8 into culture supernatant of HEK293T cells transfected with ORF8 DNA. ORF8 was probed with an anti-ORF8 antibody. (**B**) ADCC (%) mediated by plasma from 13 convalescent individuals, (**C**) 13 vaccinated individuals and (**D**) 13 previously-infected vaccinated individuals was measured using PBMCs from healthy donors as effector cells treated overnight (16 hours) with (+ORF8) or without ORF8 (-ORF8). (**E**) ADCC (%) mediated by plasmas from convalescent individuals with PBMCs or monocytes as effector cells treated overnight (+ORF8) or not (-ORF8) with ORF8. *, p < 0.05; **, p < 0.01; ***, p < 0.001; ****, p < 0.0001.

Vaccine-elicited humoral responses are associated with vaccine efficacy against SARS-CoV-2 infection and protection from severe disease (10, 11). In response to the antibodies produced by vaccinated and already infected individuals, multiple variants are emerging (31). A common feature of these variants is the apparition of multiple mutations in the Spike. These mutations decrease antibody recognition which cause immune evasion and increased transmissibility (31). Mutations were also observed in ORF8 but their effect(s) on immune evasion remain largely unknown. Our results raise the intriguing possibility that emerging variants use ORF8 to evade Fc-effector functions by impairing monocytes and to a lesser extent NK cells to mediate ADCC.

## ACKNOWLEDGMENTS

The authors thank the CRCHUM Flow Cytometry Platforms for technical assistance. This work was supported by le Ministère de l’Économie et de l’Innovation (MEI) du Québec, the Fondation du CHUM, CIHR grant nos. 352417 and 177958, and a CFI grant, 41027, to A.F, CIHR operating grant OV4-170641 to C.L.. Support was also provided by Sentinelle COVID Quebec to A.F. and C.L. A.F. is recipient of a Canada Research Chair. G.B.-B. is the recipient of an FRQS PhD fellowship. The funders had no role in study design, data collection and analysis, decision to publish, or preparation of the manuscript. We declare no competing interests.

## Material and Methods

### Ethics statement

The study was conducted in accordance with the Declaration of Helsinki in terms of informed consent and approval by an appropriate institutional board. Peripheral blood mononuclear cells (PBMCs) and plasmas from naïve, convalescent and vaccinated individuals were obtained from donors who consented to participate in this research project at CHUM. The protocol was approved by the Ethics Committee of CHUM (protocol #19.381, approved on 25 March 2020). Donors met all eligibility criteria: previously confirmed COVID-19 infection and a complete resolution of symptoms for at least 14 days.

### Cell lines and primary cells

HEK 293T human embryonic kidney cells (obtained from ATCC) were maintained at 37°C under 5% CO_2_ in Dulbecco’s Modified Eagle Medium (DMEM) (Wisent, St. Bruno, QC, Canada), supplemented with 5% fetal bovine serum (FBS) (VWR, Radnor, PA, USA) and 100 U/mL penicillin/streptomycin (Wisent). Human peripheral blood mononuclear cells (PBMCs) obtained by leukapheresis and Ficoll-Paque density gradient isolation were cryopreserved in liquid nitrogen until further use. Monocytes were isolated and separated from resting PBMC using a human CD14 positive selection kit. The selection was done according to the manufacturer protocol (EasySep™ Human CD14 Positive Selection Kit II, STEMCELL™ Technologies). PBMCs, monocytes and PBMCs depleted in mococytes were maintained in complete medium (RPMI 1640 (Thermo Fisher Scientific, Waltham, MA, USA) supplemented with 10% fetal bovine serum (FBS) (VWR, Radnor, PA, USA) and 100 U/mL penicillin/streptomycin (Wisent)). The CEM.NKr CCR5+ and CEM.NKr.Spike cells were previously described (29) and maintained in complete media.

### ORF8 Production

Three million HEK293T cells were seeded on 100 mm petri dishes. The following day, the HEK293T cells were transfected with 10 μg of ORF8 DNA using standard calcium phosphate. 16 hours later, the media was changed. 24 hours after the media was changed, the cell supernatant was centrifuged at 484 x g and aliquoted in 1.5 mL tubes. The tubes were then stored at −80°C until further use.

### ORF8 Western

45 μg lysates of HEK293T cells transfected with ORF8 DNA and 50 μl of culture supernatants were separated through 12% SDS-PAGE by electrophoresis, then transferred onto the PVDF (polyvinylidene difluoride) membrane. ORF8 protein was probed with sheep anti-ORF8 antibody (dilution 1:1000, MRC Pure Reagents, Cat. DA088) for 16 hours at 4oC, followed by incubation with donkey anti-sheep HRP (horseradish peroxidase)-conjugated secondary antibody (1:2000, Invitrogen, Cat. AP184P) for 1 hour at room temperature. The membrane was treated with the ECL (Enhanced chemiluminescence) reagent, proteins signals were exposed to the X-ray films.

### Staining of PBMCs with ORF8

Five million PBMCs were incubated with FITC-conjugated ORF8 (1 μg) in 100 μL PBS (containing 3% BSA) on ice for 30 min. The recombinant ORF8 (ThermoFisher, Cat. RP-87666) was conjugated with FITC using the FITC conjugation kit (Abcam, Cat. ab102884). Then, the anti-CD14-V450 (dilution 1:100, BD, Cat. 561390), anti-CD16-PE-Cy7 (dilution 1:100, BD, Cat. 560716) and anti-CD56-PE (dilution 1:100, BD, Cat. 556647) antibodies were added for 1 hour incubation on ice. After washing with PBS (3% BSA), cells were fixed with 4% paraformaldehyde for 15 min at room temperature, then suspended in PBS (3% BSA) and examined with LSP Fortessa flow cytometer. The data were analyzed with FlowJo.

### Bio-Layer Interferometry (BLI)

Binding kinetics were performed on an Octet RED96e system (FortéBio) at 25°C with shaking at 1,000 RPM. Amine Reactive Second-Generation (AR2G) biosensors were hydrated in water, then activated for 300 s with an S-NHS/EDC solution (Fortébio) prior to amine coupling. CD16a ectodomain (Human CD16a, amino acids Met1-Gln208 (Accession # P08637-1) with a C-terminal His-tag, ThermoFischer Scientific, catalogue number A42538) was loaded into AR2G biosensor at 12.5 μg/mL in 10mM acetate solution pH 5 (Fortébio) for 600 s and then quenched into 1M ethanolamine solution pH 8.5 (Fortébio) for 300 s. Baseline equilibration was collected for 120 s in 10X kinetics buffer. Association of ORF8 protein (SARS-CoV-2 ORF8 (aa16-121), His Tag (RP-87666), ThermoFischer Scientific) (in 10X kinetics buffer) to CD16a ectodomain was carried out for 180 s at various concentrations in a two-fold dilution series from 500nM to 31.25nM prior to dissociation for 300 s. The data were baseline subtracted prior to fitting performed using a 1:1 binding model in the FortéBio data analysis software. Calculation of on-rates (Ka), off-rates (Kdis), and affinity constants (KD) was computed using a global fit applied to all data.

### CD16 cell-surface staining on monocytes or NK cells

PBMCs, purified monocytes or PBMCs depleted in monocytes from different donors were treated with mammalian produced ORF8 for 16 hours at 37°C and 5% CO_2_. Following treatment, cells were stained with anti-CD3 BV-421, anti-CD14 PerCP-Cy5.5, anti-CD56 PE, anti-CD16 FITC and LIVE/DEAD Fixable Aqua Dead Cell Stain for 25 minutes at 4°C to identify the monocytes and NK cells among the PBMC population and to measure their surface level of CD16. After staining, the cells were fixed with 2% PFA and stored at 4°C until the samples were acquired on a LSRII cytometer (BD Biosciences, Mississauga, ON, Canada). Data analysis was performed using FlowJo v10.5.3 (Tree Star, Ashland, OR, USA).

### Antibody-dependent cellular cytotoxicity (ADCC) assay

This assay was previously described (29, 30). Briefly, for evaluation of anti-SARS-CoV-2 ADCC activity, parental CEM.NKr CCR5+ cells were mixed at a 1:1 ratio with CEM.NKr-Spike cells. These cells were stained for viability (AquaVivid; Thermo Fisher Scientific) and a cellular dye (cell proliferation dye eFluor670; Thermo Fisher Scientific) and subsequently used as target cells. PBMCs, purified monocytes or PBMCs depleted in monocytes were treated or not with ORF8 (which was produced in mammalian cells) overnight for 16 hours. These cells were stained with another cellular marker (cell proliferation dye eFluor450; Thermo Fisher Scientific) and used as effector cells. Stained effector and target cells were mixed at a 10:1 ratio in 96-well V-bottom plates. Plasma from convalescent, vaccinated or convalescent and vaccinated individuals at a dilution of 1/500 was added to the appropriate wells. The plates were subsequently centrifuged for 1 min at 300 x g, and incubated at 37°C, 5% CO_2_ for 5 h before being fixed in a 2% PBS-formaldehyde solution. Since CEM.NKr-Spike cells express GFP, ADCC activity was calculated using the formula: [(% of GFP + cells in Targets plus Effectors) - (% of GFP + cells in Targets plus Effectors plus plasma)]/(% of GFP + cells in Targets) x 100 by gating on transduced live target cells. All samples were acquired on an LSRII cytometer (BD Biosciences, Mississauga, ON, Canada) and data analysis performed using FlowJo v10.5.3 (Tree Star, Ashland, OR, USA).

### Statistical analyses

Statistics were analyzed using GraphPad Prism version 8.0.2 (GraphPad, San Diego, CA, (USA). Every data set was tested for statistical normality and this information was used to apply the appropriate (parametric or nonparametric) statistical test. P values <0.05 were considered significant; significance values are indicated as * P<0.05, ** P<0.01, *** P<0.001, **** P<0.0001.

---

**Supplemental Figure 1.**
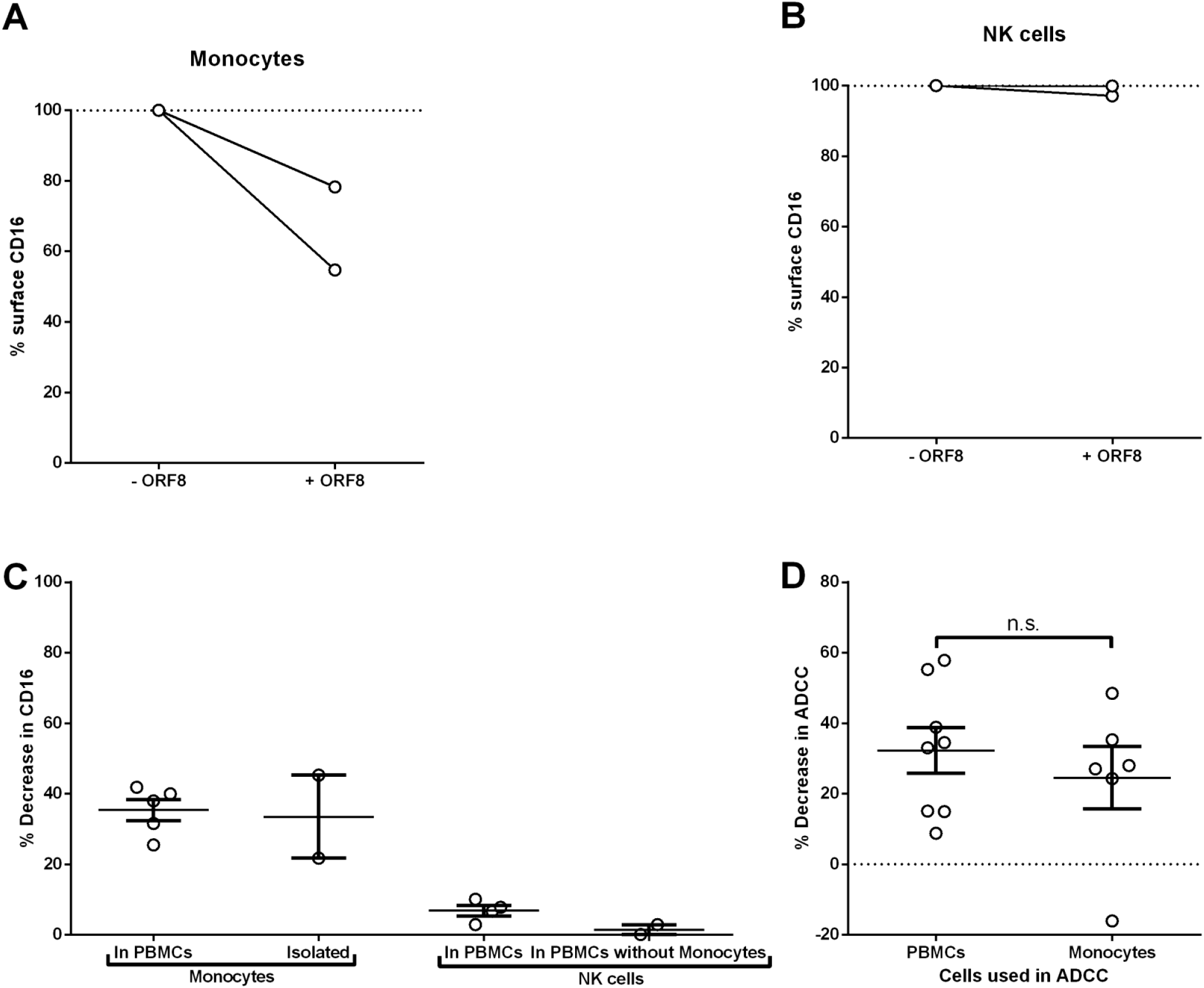
ORF8 modulation of CD16 levels at the surface of purified monocytes and NK cells. (**A**) Purified monocytes from PBMCs were treated (16h) or not with ORF8 and cell surface levels of CD16 were measured by flow cytometry, n=2. (**B**) Monocytes-depleted PBMCs were treated (16h) or not with ORF8 and CD16 levels were measured at the surface of NK cells, n=2. (**C**) Decrease of surface CD16 after ORF8 treatment on monocytes present in the PBMC population, on isolated monocytes, on NK cells in the PBMC population and on NK cells in monocytes-depleted PBMC population. (**D**) Decrease of ADCC mediated by PBMCs or purified monocytes after treatment with ORF8 for 16 hours. Statistical significance was evaluated using a parametric t-test for (D). n.s., not significant. (C) and (D) Mean values ± the standard error of the mean (SEM).

## References

1. Huang C, Wang Y, Li X, Ren L, Zhao J, Hu Y, Zhang L, Fan G, Xu J, Gu X, Cheng Z, Yu T, Xia J, Wei Y, Wu W, Xie X, Yin W, Li H, Liu M, Xiao Y, Gao H, Guo L, Xie J, Wang G, Jiang R, Gao Z, Jin Q, Wang J, Cao B. 2020. Clinical features of patients infected with 2019 novel coronavirus in Wuhan, China. Lancet 395:497–506.

2. Wu F, Zhao S, Yu B, Chen YM, Wang W, Song ZG, Hu Y, Tao ZW, Tian JH, Pei YY, Yuan ML, Zhang YL, Dai FH, Liu Y, Wang QM, Zheng JJ, Xu L, Holmes EC, Zhang YZ. 2020. A new coronavirus associated with human respiratory disease in China. Nature 579:265–269.

3. Zhu N, Zhang D, Wang W, Li X, Yang B, Song J, Zhao X, Huang B, Shi W, Lu R, Niu P, Zhan F, Ma X, Wang D, Xu W, Wu G, Gao GF, Tan W, China Novel Coronavirus I, Research T. 2020. A Novel Coronavirus from Patients with Pneumonia in China, 2019. N Engl J Med 382:727–733.

4. Finkel Y, Mizrahi O, Nachshon A, Weingarten-Gabbay S, Morgenstern D, Yahalom-Ronen Y, Tamir H, Achdout H, Stein D, Israeli O, Beth-Din A, Melamed S, Weiss S, Israely T, Paran N, Schwartz M, Stern-Ginossar N. 2021. The coding capacity of SARS-CoV-2. Nature 589:125–130.

5. Tang X, Wu C, Li X, Song Y, Yao X, Wu X, Duan Y, Zhang H, Wang Y, Qian Z, Cui J, Lu J. 2020. On the origin and continuing evolution of SARS-CoV-2. Natl Sci Rev 7:1012–1023.

6. Flower TG, Buffalo CZ, Hooy RM, Allaire M, Ren X, Hurley JH. 2021. Structure of SARS-CoV-2 ORF8, a rapidly evolving immune evasion protein. Proc Natl Acad Sci U S A 118.

7. Zhang Y, Chen Y, Li Y, Huang F, Luo B, Yuan Y, Xia B, Ma X, Yang T, Yu F, Liu J, Liu B, Song Z, Chen J, Yan S, Wu L, Pan T, Zhang X, Li R, Huang W, He X, Xiao F, Zhang J, Zhang H. 2021. The ORF8 protein of SARS-CoV-2 mediates immune evasion through down-regulating MHC-Iota. Proc Natl Acad Sci U S A 118.

8. Wu X, Manske MK, Ruan G, Nowakowski KE, Abeykoon JP, Tang X, Yu Y, Witter TL, Taupin V, Paludo J, Ansell SM, Badley AD, Schellenberg MJ, Witzig TE. 2021. Secreted ORF8 is a pathogenic cause of severe Covid-19 and potentially targetable with select NLRP3 inhibitors. bioRxiv.

9. Lin X, Fu B, Yin S, Li Z, Liu H, Zhang H, Xing N, Wang Y, Xue W, Xiong Y, Zhang S, Zhao Q, Xu S, Zhang J, Wang P, Nian W, Wang X, Wu H. 2021. ORF8 contributes to cytokine storm during SARS-CoV-2 infection by activating IL-17 pathway. iScience 24:102293.

10. Khoury DS, Cromer D, Reynaldi A, Schlub TE, Wheatley AK, Juno JA, Subbarao K, Kent SJ, Triccas JA, Davenport MP. 2021. Neutralizing antibody levels are highly predictive of immune protection from symptomatic SARS-CoV-2 infection. Nat Med 27:1205–1211.

11. Earle KA, Ambrosino DM, Fiore-Gartland A, Goldblatt D, Gilbert PB, Siber GR, Dull P, Plotkin SA. 2021. Evidence for antibody as a protective correlate for COVID-19 vaccines. Vaccine 39:4423–4428.

12. Ullah I, Prevost J, Ladinsky MS, Stone H, Lu M, Anand SP, Beaudoin-Bussieres G, Symmes K, Benlarbi M, Ding S, Gasser R, Fink C, Chen Y, Tauzin A, Goyette G, Bourassa C, Medjahed H, Mack M, Chung K, Wilen CB, Dekaban GA, Dikeakos JD, Bruce EA, Kaufmann DE, Stamatatos L, McGuire AT, Richard J, Pazgier M, Bjorkman PJ, Mothes W, Finzi A, Kumar P, Uchil PD. 2021. Live imaging of SARS-CoV-2 infection in mice reveals that neutralizing antibodies require Fc function for optimal efficacy. Immunity 54:2143–2158 e15.

13. Beaudoin-Bussieres G, Chen Y, Ullah I, Prevost J, Tolbert WD, Symmes K, Ding S, Benlarbi M, Gong SY, Tauzin A, Gasser R, Chatterjee D, Vezina D, Goyette G, Richard J, Zhou F, Stamatatos L, McGuire AT, Charest H, Roger M, Pozharski E, Kumar P, Mothes W, Uchil PD, Pazgier M, Finzi A. 2022. A Fc-enhanced NTD-binding non-neutralizing antibody delays virus spread and synergizes with a nAb to protect mice from lethal SARS-CoV-2 infection. Cell Rep 38:110368.

14. Brunet-Ratnasingham E, Anand SP, Gantner P, Dyachenko A, Moquin-Beaudry G, Brassard N, Beaudoin-Bussieres G, Pagliuzza A, Gasser R, Benlarbi M, Point F, Prevost J, Laumaea A, Niessl J, Nayrac M, Sannier G, Orban C, Messier-Peet M, Butler-Laporte G, Morrison DR, Zhou S, Nakanishi T, Boutin M, Descoteaux-Dinelle J, Gendron-Lepage G, Goyette G, Bourassa C, Medjahed H, Laurent L, Rebillard RM, Richard J, Dube M, Fromentin R, Arbour N, Prat A, Larochelle C, Durand M, Richards JB, Chasse M, Tetreault M, Chomont N, Finzi A, Kaufmann DE. 2021. Integrated immunovirological profiling validates plasma SARS-CoV-2 RNA as an early predictor of COVID-19 mortality. Sci Adv 7:eabj5629.

15. Zohar T, Loos C, Fischinger S, Atyeo C, Wang C, Slein MD, Burke J, Yu J, Feldman J, Hauser BM, Caradonna T, Schmidt AG, Cai Y, Streeck H, Ryan ET, Barouch DH, Charles RC, Lauffenburger DA, Alter G. 2020. Compromised Humoral Functional Evolution Tracks with SARS-CoV-2 Mortality. Cell 183:1508–1519 e12.

16. Keeler SP, Fox JM. 2021. Requirement of Fc-Fc Gamma Receptor Interaction for Antibody-Based Protection against Emerging Virus Infections. Viruses 13.

17. Corrales-Aguilar E, Trilling M, Hunold K, Fiedler M, Le VT, Reinhard H, Ehrhardt K, Merce-Maldonado E, Aliyev E, Zimmermann A, Johnson DC, Hengel H. 2014. Human cytomegalovirus Fcgamma binding proteins gp34 and gp68 antagonize Fcgamma receptors I, II and III. PLoS Pathog 10:e1004131.

18. Gu W, Guo L, Li R, Niu J, Luo X, Zhang J, Xu Y, Tian Z, Feng L, Wang Y. 2016. Elevated plasma-soluble CD16 levels in porcine reproductive and respiratory syndrome virus-infected pigs: correlation with ADAM17-mediated shedding. J Gen Virol 97:632–638.

19. Oliviero B, Mantovani S, Varchetta S, Mele D, Grossi G, Ludovisi S, Nuti E, Rossello A, Mondelli MU. 2017. Hepatitis C virus-induced NK cell activation causes metzincin-mediated CD16 cleavage and impaired antibody-dependent cytotoxicity. J Hepatol 66:1130–1137.

20. Anand SP, Finzi A. 2019. Understudied Factors Influencing Fc-Mediated Immune Responses against Viral Infections. Vaccines (Basel) 7.

21. Veillette M, Richard J, Pazgier M, Lewis GK, Parsons MS, Finzi A. 2016. Role of HIV-1 Envelope Glycoproteins Conformation and Accessory Proteins on ADCC Responses. Curr HIV Res 14:9–23.

22. Veillette M, Desormeaux A, Medjahed H, Gharsallah NE, Coutu M, Baalwa J, Guan Y, Lewis G, Ferrari G, Hahn BH, Haynes BF, Robinson JE, Kaufmann DE, Bonsignori M, Sodroski J, Finzi A. 2014. Interaction with cellular CD4 exposes HIV-1 envelope epitopes targeted by antibody-dependent cell-mediated cytotoxicity. J Virol 88:2633–44.

23. Veillette M, Coutu M, Richard J, Batraville LA, Dagher O, Bernard N, Tremblay C, Kaufmann DE, Roger M, Finzi A. 2015. The HIV-1 gp120 CD4-Bound Conformation Is Preferentially Targeted by Antibody-Dependent Cellular Cytotoxicity-Mediating Antibodies in Sera from HIV-1-Infected Individuals. J Virol 89:545–51.

24. Prevost J, Richard J, Medjahed H, Alexander A, Jones J, Kappes JC, Ochsenbauer C, Finzi A. 2018. Incomplete Downregulation of CD4 Expression Affects HIV-1 Env Conformation and Antibody-Dependent Cellular Cytotoxicity Responses. J Virol 92.

25. Costa J, Yee C, Nakamura Y, Rabson A. 1978. Characteristics of the Fc receptor induced by herpes simplex virus. Intervirology 10:32–9.

26. Johnson DC, Feenstra V. 1987. Identification of a novel herpes simplex virus type 1-induced glycoprotein which complexes with gE and binds immunoglobulin. J Virol 61:2208–16.

27. Litwin V, Sandor M, Grose C. 1990. Cell surface expression of the varicella-zoster virus glycoproteins and Fc receptor. Virology 178:263–72.

28. Thale R, Lucin P, Schneider K, Eggers M, Koszinowski UH. 1994. Identification and expression of a murine cytomegalovirus early gene coding for an Fc receptor. J Virol 68:7757–65.

29. Beaudoin-Bussieres G, Richard J, Prevost J, Goyette G, Finzi A. 2021. A new flow cytometry assay to measure antibody-dependent cellular cytotoxicity against SARS-CoV-2 Spike-expressing cells. STAR Protoc 2:100851.

30. Anand SP, Prevost J, Nayrac M, Beaudoin-Bussieres G, Benlarbi M, Gasser R, Brassard N, Laumaea A, Gong SY, Bourassa C, Brunet-Ratnasingham E, Medjahed H, Gendron-Lepage G, Goyette G, Gokool L, Morrisseau C, Begin P, Martel-Laferriere V, Tremblay C, Richard J, Bazin R, Duerr R, Kaufmann DE, Finzi A. 2021. Longitudinal analysis of humoral immunity against SARS-CoV-2 Spike in convalescent individuals up to 8 months post-symptom onset. Cell Rep Med 2:100290.

31. Prevost J, Finzi A. 2021. The great escape? SARS-CoV-2 variants evading neutralizing responses. Cell Host Microbe 29:322–324.

